# Absence of pathogenic Short Tandem Repeat expansions in Systemic Lupus Erythematosus disease-associated genes

**DOI:** 10.1101/729467

**Authors:** Audrey Lee, Vicky Cho, T. Daniel Andrews

**Affiliations:** Genome Informatics Laboratory, Department of Immunology and Infectious Disease, The John Curtin School of Medical Research, The Australian National University, ACT Australia

## Abstract

Short tandem repeat (STR) expansions have been shown to be pathogenic in human neurological diseases, such as Huntington disease. Yet, the potential role of STRs in non-neurological diseases has yet to be fully investigated. In this study, the potential role of STR expansions in the pathogenesis of systemic lupus erythematosus (SLE) was investigated using patient genomic data and two computational tools, HipSTR and exSTRa. The length variability of STRs in 76 SLE-associated genes was compared using exome data from 271 SLE affected individuals and 158 of their unaffected relatives. We conclude that no large STR expansions associated with SLE were present in these affected individuals within the 76 genes investigated. Lack of evidence does not negate a pathogenic role for STR expansions in SLE, yet given the number of individuals included in this study, we expect that this is not a common source of pathogenesis in SLE.

**Significance statement:** The increasing availability and decreasing cost of sequencing genomes lends itself to computational analysis, extracting information to aid diagnosis, guide treatment or discover disease mechanisms and new treatments. Computational tools have been developed to look for various types of mutations, including short tandem repeats (STRs), which has been shown to cause diseases such as Huntington disease. Limited research on the possible role of STR expansions in systemic lupus erythematosus (SLE) has been done. Here we use computational tools to compare the length of STRs in 76 SLE-associated genes in patients and their unaffected relatives. Our results did not identify any large STR expansions associated with SLE, and further research is required to gain a better understanding of this complex disease.

## Introduction

Short tandem repeats (STRs) are repeating units of 2-6 basepairs found in genomic DNA (1). Since the early 1990s, changes to the length of STRs have been shown to cause disease, particularly in the field of neurological disease, e.g. Huntington disease and fragile X syndrome (2). One mechanism by which STRs can cause disease is by extending the amino acid sequences in expressed proteins. This has been shown in Huntington disease, where the trinucleotide CAG repeat expansion encodes the Huntingtin protein with a pathological extended polyglutamine tract (3). More recently, the molecular and cellular dysfunction caused by STRs have also been shown to be a major genetic contribution to other diseases, such as frontotemporal dementia and amyotrophic lateral sclerosis (4). With the abundance of genetic sequencing information and tools available, there is growing interest to elucidate if STRs are responsible for the genetic component of other non-neurological diseases.

Autoimmune diseases have a multifactorial aetiology, including both environmental and genetic contributions (5). Systemic lupus erythematosus (SLE) is considered to be a prototypic autoimmune disease and is characterised by the production of anti-nuclear antibodies with a heterogenous clinical presentation. The aetiology of SLE is not well understood, although strong familial aggregation supports the involvement of genetic factors. Twin studies in SLE have shown a concordance of approximately 5% in dizygotic twins, and 25-50% in identical twins (6).

While several single nucleotide polymorphisms (SNPs) have already been identified and functionally characterised for pathogenic contribution to SLE (7, 8), the investigation of STRs in the genetic component of SLE has been limited. Prior to the advent of next-generation sequencing and analysis tools, STR detection methods were limited to polymerase chain reactions (PCR) or Southern blots. These methods are costly in time and resources, and cannot easily be upscaled to analyse multiple loci, but they remain to be the current gold standard (9). Previous work in SLE by these methods have identified an association between the Fli1 promoter STR length and the absence of nephritis (10), as well as identifying STRs as a marker for disease susceptibility in HLA Class III (11).

The identification of STRs has been made easier by next-generation sequencing, which provides access to whole genome or exome sequences. Unlike traditional methods, the computational analysis of these sequences can identify multiple different repeat expansions in a shorter period of time, and hence can be useful in diagnosis (9). Furthermore, when used in conjunction with disease status or presentation, the segregation of STR lengths can uncover new pathogenic STRs, furthering the understanding of disease mechanisms.

Tools to facilitate the analysis of STRs in exome and genome data have been emerging, with newer tools able to identify longer STRs, and with greater confidence. One of these methods is HipSTR (12), a tool that uses an expectation maximisation algorithm to report STR alleles and length. HipSTR is constrained to identifying alleles with repeat length shorter than the sequencing read length, which is typically 150 basepairs in short-read sequencing (13). This may be an obstacle in the identification of possibly pathogenic STRs, as it has been suggested that pathogenic STR mutations are typically long (exceeding 100 basepairs), which has been observed in Mendelian human diseases (9).

Tools such as exSTRa (14) overcome this issue by determining the repeat content in all reads that contain STRs, whether in full or in partial, and hence are not limited to STRs shorter than 150 basepairs. Read fragments that are contained entirely within expanded alleles are excluded by exSTRa from analysis in order to prevent any ambiguous mapping (13), unlike tools such as ExpansionHunter (15) and TREDPARSE (16).

The dataset used was from the Australian Point Mutation in Systemic Lupus Erythematosus (APOSLE) study, which has also been involved in publications on immunological effector molecules and SNPs in SLE (17, 18). The exomes of 271 individuals diagnosed with SLE and 158 unaffected relatives from the APOSLE study were analysed using HipSTR and exSTRa to look for lengthened STRs in 76 SLE-associated genes. Our findings suggest that there are no large STR expansions in SLE-associated loci contributing to pathogenicity in SLE.

## Results

Two similar computational tools were applied to our SLE cohort data to reconstruct the STR regions associated with a set of putative SLE-linked disease genes. These 76 SLE genes were collated from multiple sources by Jiang et al., encompassing all identified monogenic causes of SLE, all known causes of interferonopathies and other genes linked with SLE via GWAS evidence (19).

### HipSTR analysis

Analysis of SLE cohort genome data with the HipSTR tool identified 14 STRs that were expanded in SLE-affected individuals of the APOSLE cohort (Table 1). These STRs exceeded the maximum repeat length found in unaffected relatives in the APOSLE cohort, and in two additional healthy references: MGRB (20) and STR Catalog (21). The expansions were modest, ranging from 1-5 additional motifs. Expanded STRs were observed in unrelated individuals and also in familial clusters, with the most prevalent STR present in 6 affected individuals. However, no intrafamilial segregation of disease status with mutation was observed, either due to lack of occurrence or incomplete genetic information from relatives. A complete table of all STRs analysed by HipSTR is provided in supplementary information (Table S1).

**Table 1.** Expanded short tandem repeats (STRs) in SLE-affected individuals from APOSLE cohort.

| *Gene* | *Repeat motif* | *Start hg19* | *Number of individuals* | *Expanded repeat length* | *APOSLE unaffected maximum repeat* | *MGRB maximum repeat* | *STR Catalog maximum repeat* |
| --- | --- | --- | --- | --- | --- | --- | --- |
| *ARID1A* | GCC | 27023372 | 1 | 9.3 | 8.3 | -<sup>a</sup> | 8.0 |
| *ARID1B* | CAG | 157099403 | 1 | 20.7 | 19.7 | -<sup>a</sup> | 19.3 |
| *CEP164* | CCT | 117280716 | 1 | 8.0 | 7.0 | 7.0 | 7.0 |
| *HLX* | CAG | 221053581 | 2 | 15.3 | 10.3 | 10.3 | 11.0 |
| *IL27* | TCC | 28511172 | 2 | 16.7 | 14.7 | 14.7 | 14.3 |
| *KRT10* | CTC | 38978747 | 1 | 15.3 | 14.3 | 14.3 | 15.0 |
| *NKX2-1* | CCG | 36986880 | 6 | 10.3 | 7.3 | 7.3 | 7.3 |
| *PCSK9* | CTG | 55505553 | 1 | 9.7 | 8.7 | 8.7 | 8.3 |
| *POLG* | GCT | 89876820 | 1 | 18.7 | 15.7 | 14.7 | 15.3 |
| *POMC* | GCT | 25384460 | 1 | 13.3 | 10.3 | 10.3 | 10.0 |
| *SFTPC* | GTG | 22020159 | 1 | 6.7 | 5.7 | 5.7 | 6.3 |
| *SLC19A2* | GCC | 169454958 | 1 | 8.7 | 7.7 | 7.7 | 7.3 |
| *TGM1* | TCTGGC | 24731464 | 4 | 4.8 | 3.8 | 3.8 | 3.7 |
| *ZMIZ1* | CTC | 81070787 | 1 | 6.7 | 5.7 | 5.7 | 5.3 |

<sup>a</sup>MGRB data was not available for these STRs.

### exSTRa analysis

The algorithmic approach in exSTRa differs from that in HipSTR – exSTRa determines the repeat length from each read sequence directly while HipSTR uses alignment methods. exSTRa looks for the proportion of the read showing the repeat motif at any starting base and does not require the repeat to be continuous. From this, a repeat score for each read is generated, which equals the number of bases that are part of the repeat motif in a read. An empirical cumulative distribution function (ECDF) plot for the repeat scores of each locus were plotted – two of the 216 plots generated from the loci analysed have been provided for discussion (Figure 1). In Figure 1, poor read depth is observed in some samples, forming almost vertical lines due to lack of stepping in the graph.

**Figure 1:**
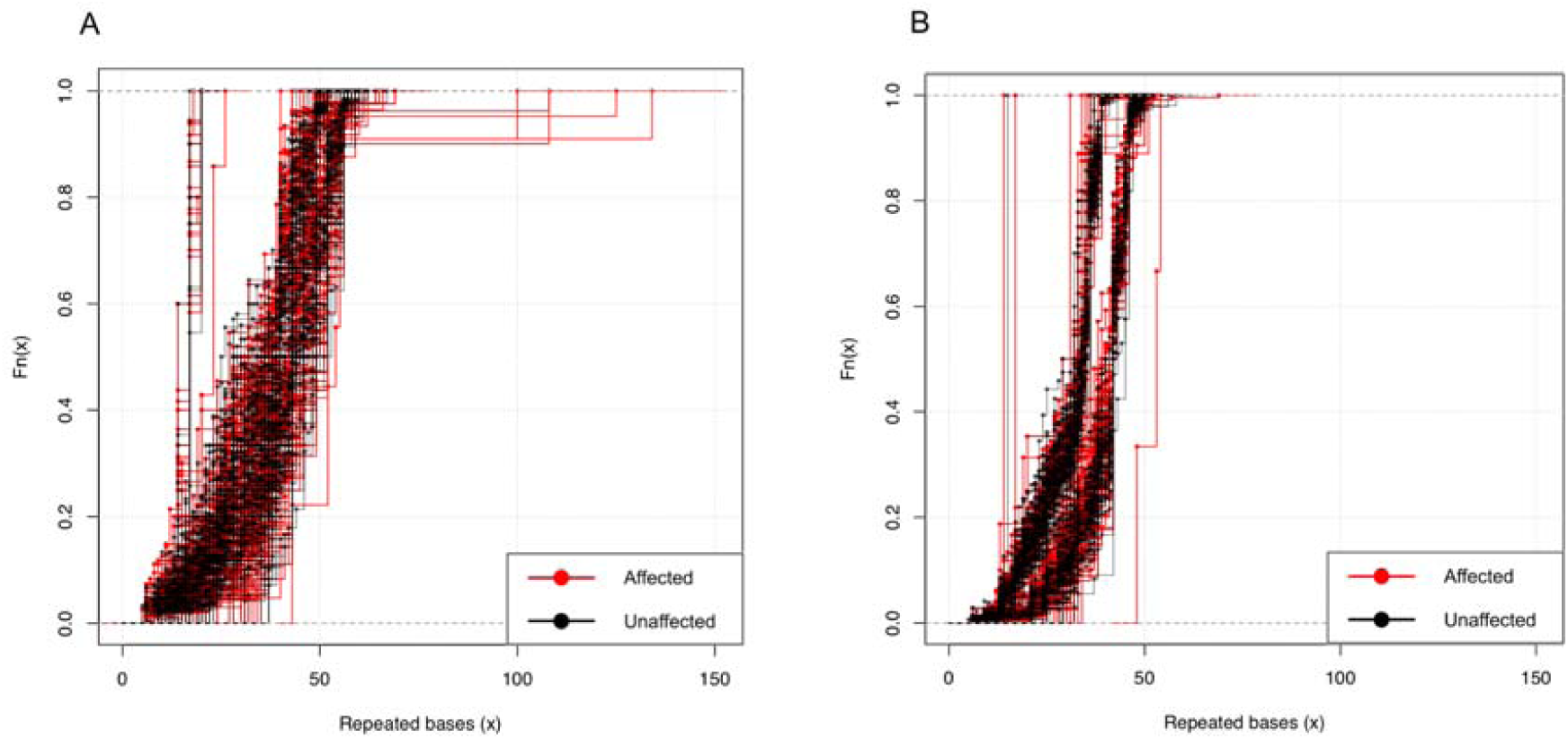
exSTRa ECDF plots of repeat scores from analysis of affected and unaffected individuals in APOSLE cohort: a) STR (CTC repeat) on KRT10 (hg19 start: 38978747) start; b) STR (GCA repeat) on NBEAL2 (hg19 start: 47043575)

The 216 ECDF plots were visually analysed according to exSTRa documentation, and no STR expansions were found. Additional ECDF plots corresponding to the expanded STRs identified by HipSTR in Table 1 have been provided in supplementary information (Figure S1). Expanded samples have more reads with higher repeat scores, and hence are recognised by a shift of data points to the right. For dominant disorders, we would expect to find this right shift in the upper half of the plot, as approximately half of the reads would have higher than average repeat scores. For recessive disorders, we would expect to find this right shift for all reads. As an example, Figure 1a shows a right shift for approximately the top 10% of the plot and within this, it appears only one read accounts for the extreme right shift. This is unlikely to be a true expansion.

The principle behind exSTRa is an outlier detection method, which aims to classify each sample as normal or expanded. At each locus, underestimations can result from mutations and impure repeats, and overestimates can result from motif matching outside the STR. However, these inaccuracies at a locus should be systematically biased, and hence consistent for all samples at the same locus. While this can be used to compare between cases and controls, we cannot expect an accurate numerical STR length from this method. This accounts for the discrepancy between the maximum lengths observed between HipSTR and exSTRa, such as for the KRT10 STR, which was 15.3 repeats (46 basepairs) in HipSTR but longer in exSTRa (Figure 1a).

Variation within the population is sometimes observed, with two groups of different lengths becoming distinct. An example given is the STR on NBEAL2 (Figure 1b), where there is a distinct lack of segregation between affected and unaffected samples, represented in red and black respectively. Segregation between affected and unaffected samples was not observed within any of the 216 plots from each of the loci analysed.

## Discussion

HipSTR identified 14 STRs which were longer in certain affected individuals than in any of the control groups. While this would align with what is expected for pathogenic mutations, the repeats are not very much longer – the largest repeat of 4.3 was for the CAG repeat on HLX, where the affected maximum was 15.3 repeats and the control maximum was 11. From observations in Mendelian human diseases, it has been suggested that pathogenic STR mutations are typically long (exceeding 100 basepairs) (9). Hence, it is unlikely that these 14 STRs identified by HipSTR are unambiguously pathogenic – and so at this point in time, it would be difficult to justify further focused research on these STRs until additional evidence surfaces. It is important to note that the STR-read length limitation which is intrinsic to HipSTR does not apply to exSTRa. No expanded STRs were identified by exSTRa, which supports the suggestion that the 14 expanded STRs from HipSTRs are unlikely to be pathogenic. While certain samples did show insufficient read depth on the ECDF plots, the exomes used had a median read depth coverage of 174.9 – hence, most samples would have had sufficient read depth at most loci.

A limitation of the analysis in this study is that no positive controls were available to test the reliability of the tools used. Both HipSTR and exSTRa reference the involvement of positive controls in their respective papers. The HipSTR publication benchmarked the results from their tool analysis against gold-standard capillary electrophoresis, and also outperformed other tools available for STR analysis at the time (12). The exSTRa publication tested and proved performance with a simulation study involving four retrospective cohorts with eleven known repeat-expansion disorders, which included dominant, recessive and X-linked disorders (14). These tools show promising ability to identify expanded STRs. However, the genomes used as positive controls are not made available to the public for obvious ethical and privacy concerns. Without this access, it is difficult to establish a positive control in our study, and we are necessitated to rely on previously reported results. The identification of any STR expansions, SLE-related or not, within datasets accessible to us would facilitate the replication of positive findings as a control for future studies.

The control used in this study for both HipSTR and exSTRa were the unaffected relatives of the SLE-diagnosed relatives. As a result, these controls all have a family history of SLE. Benefits of this approach include the ability to identify the boundary between affected and unaffected, had a pathological STR expansion been present. Similarly, genetic anticipation may have been observed in related individuals. However, there are also limitations of using unaffected relatives as controls. Under the common variant common disease hypothesis, a risk-increasing mutation is likely to be present with relatives of the affected, with an extra external factor tipping the balance towards disease (5). In such a situation, this intrafamilial comparison will not highlight these risk-increasing variants. While comparison with MGRB and STR Catalog for the HipSTR data filled this gap and provided control data of unaffected individuals without a family history, there was no access to the equivalent for exSTRa data. Future studies would benefit from access to exome sequences of unaffected individuals without a family history.

While the results from this study suggest that STRs were not found to contribute to pathogenicity in these SLE individuals, there are still other avenues to explore before ruling out the possibility. Exome data has been used in this study, and has been shown to be valid for both HipSTR (12) and exSTRa (14). However, STRs are not limited to protein dysfunction, and non-coding repeat expansions have been associated with disease, for example, FRDA (3). It has been suggested that STRs may have a regulatory role: affecting transcription factor binding, promotor distances and forming irregular DNA structures (1). Hence, it would be reasonable to investigate STRs in genome data, particularly as whole genome sequencing becomes more affordable.

The 76 genes associated with SLE that were included in this study were based on a paper by Jiang et al. (21). However, another study has suggested that a set of 98 genes may be a better representation of the genes associated with SLE (24). Alternatively, given sufficient computational resources, it would also be possible to search across the genome for expansions at all STRs; using a database such as STR Catalog (23) as a guide.

Tools to identify STRs in genomic data have emerged and progressed rapidly over the last few years, dealing with challenges such as misalignments, stutter errors and read length (13). There is still room for improvement, with the need of tools that provide numerical answers with confidence and without read length limitations. These would be useful and easy to implement into pipelines for diagnosis. The emergence of long-read sequencing is also an exciting addition to the field and will likely aid the discovery of more pathogenic STRs in the future.

## Materials and methods

### Dataset

Illumina whole exome sequencing data for 429 individuals was analysed, with 271 SLE-affected individuals and 158 unaffected relatives. These trios or multiplex families were recruited and genotyped as part of the APOSLE (Australian Point Mutation in Systemic Lupus Erythematosus) study, for which ethics approval was obtained from each participating institution (17, 18). 2 SLE-affected individuals were excluded from the exSTRa analysis due to technical incompatibilities. Prior to analysis for pathogenic STR expansions, all exomes had been assessed for plausible single nucleotide variants by which to attribute disease pathogenicity (19, 20). In all cases pathogenicity could not be explained by known, monogenic sequence variation or false-positive variant miscalls.

### Regions

76 genes associated with SLE were provided by Jiang et al. (21). STRs within these associated genes were identified for analysis using the STR Catalog database (22), resulting in a total of 219 SLE-associated STRs. Final analysis was conducted on 216 STRs, as the following three were excluded due to technical incompatibilities: ARID1A GCG repeat (hg19 start: 27023912), ARFIP2 GCA repeat (hg19 start: 6502755) and ANKRD17 GCC repeat (hg19 start: 74124056).

### HipSTR

Each individual from the APOSLE study had a corresponding sequence alignment data in a BAM file format. For some of these files, HipSTR produced an error and was unable to analyse them. We identified a double LB tag artefact in the header which needed to be removed in order for the error to resolve. Samtools (25) was used to abstract the header as a text file from each of the 129 BAM files with a double LB tag. A script was written in Shell which used pattern matching to remove one copy of the LB tag used, producing a new text file which was then to attached to the BAM files by Samtools using the reheader command. A new BAM index file was then created for each BAM file due to the changes made.

A script for HipSTR was written including the 429 BAM files, a hs37d5 alignment, a regions file (see above Regions) and the pre-set 100 minimum reads. This script was submitted to the Raijin supercomputer which is part of the National Computational Infrastructure (NCI), requiring 1.56 service units, 6.45 GB of memory and 1834 seconds of CPU time. Call filtering was performed on the output from HipSTR, using the *script filter_vcf.py* which is included in the HipSTR package. The default settings were used: min-call-qual 0.9, max-call-flank-indel 0.15, max-call-stutter 0.15, min-call-allele-bias −2 and min-call-strand-bias −2.

The filtered output was analysed by a script which used the VariantAnnotation package (26) in R (27). STRs with more than 20% of genotypes missing were removed from analysis, as were STRs that had the reference genotype in all samples. Analysis was centred on data extracted from the GB heading in the output file, which corresponded to the base pair differences of each genotype from the reference. This was converted to a number of repeats by dividing the base pair difference by the motif length and adding the result to the reference number of repeats. The maximum repeat length was identified for the affected and unaffected populations. Initial screening was performed to identify STRs which had a higher maximum in affected population compared to the unaffected population.

Further comparison was performed with the maximum repeat length observed in two external datasets: MGRB (20) and STR Catalog (23). MGRB (Medical Genome Reference Bank) (22) is a genome database of 4000 healthy older individuals from Australia. These genomes were analysed using HipSTR to identify the normal length of STRs in SLE-associated genes. The MGRB database did not have information for some STRs and this has been indicated for the relevant STRs in Table 1 and Table S1. STR Catalog (23) is an STR database of over 1000 individuals from the 1000 Genomes Project. All data comparisons were performed using scripts written in R, including the final identification of STRs which were longer in the affected APOSLE cohort than all three normal references (unaffected APOSLE cohort, MGRB, and STR Catalog). For these expanded STRs, the familial connections of the individuals involved were examined using unidentified patient information.

### exSTRa

Due to technical incompatibilities, two BAM files were removed from exSTRa analysis resulting in the analysis of 269 affected and 158 unaffected individuals. A regions file was created for exSTRa, by manually editing the regions file used for HipSTR. This regions file included the name, chromosome number, hg19 start, end and motif for each of the 216 STRs used in the HipSTR analysis. An exSTRa program script for each of the 427 BAM files was written using a shell template and the BAM file locations, and these were then submitted to NCI, using up to 0.33 service units and 830 seconds of CPU time to process each script. The script for each BAM file returned individual output files which were combined and analysed using the exSTRa package in R, with additional input to specify which individuals were affected or unaffected. An empirical cumulative distribution function (ECDF) was then plotted for each locus, using different colours to identify affected and unaffected individuals. Visual analysis of the 216 plots was performed manually, looking for distinctive features of a potentially pathogenic expanded STR as outlined in exSTRa documentation (14).

## Acknowledgments

We grateful to Melanie Bahlo and Mark Bennett for discussions and guidance with the application of exSTRa to exome data. We thank the Medical Genome Reference Bank for granting full access to sequence data of health control individuals. Exome sequencing from patient material was provided by the Centre for Personalised Immunology, an Australian NHMRC centre of research excellence, based within the John Curtin School of Medical Research. This project was undertaken with the resources of the National Computational Infrastructure (NCI), which is supported by the Australian Government.

