## Supplementary Table S1 for "Absence of pathogenic Short Tandem Repeat expansions in Systemic Lupus Erythematosus disease-associated genes"

**Table S1. HipSTR analysis of short tandem repeat (STRs) in SLE-affected individuals from APOSLE cohort, in comparison to three healthy controls: unaffected relatives, MGRB (1) and STR Catalog (2). Chr, chromosome.**

| Gene | Chr | Repeat motif | Start hg19 | End hg19 | Expanded repeat length | APOSLE unaffected maximum repeat | MGRB maximum repeat | STR Catalog maximum repeat |
| --- | --- | --- | --- | --- | --- | --- | --- | --- |
| ACADVL | 17 | CCCGG | 7123470 | 7123490 | 4.20 | 4.20 | 4.20 | 4.20 |
| ADAMTS13 | 9 | CAGAGG | 136289464 | 136289488 | 4.17 | 4.17 | 4.17 | 4.00 |
| AGPAT2 | 9 | CAG | 139581759 | 139581780 | 8.33 | 8.33 | 8.33 | 8.00 |
| ANKRD17 | 4 | CGC | 74124271 | 74124295 | 8.33 | 8.33 | 9.00 | 10.00 |
| ARID1A | 1 | GGC | 27023141 | 27023172 | 10.67 | 10.67 | n/a | 10.33 |
| ARID1A | 1 | CGG | 27023257 | 27023275 | 6.33 | 6.33 | n/a | 6.00 |
| ARID1A | 1 | GCC | 27023372 | 27023396 | 9.33 | 8.33 | n/a | 8.00 |
| ARID1A | 1 | GCA | 27100182 | 27100205 | 9.00 | 9.00 | 9.00 | 9.67 |
| ARID1A | 1 | GAA | 27105676 | 27105690 | 5.00 | 5.00 | 5.00 | 4.67 |
| ARID1B | 6 | TCC | 157099166 | 157099186 | 7.00 | 9.00 | 7.00 | 7.00 |
| ARID1B | 6 | CCA | 157099303 | 157099375 | 26.33 | 26.33 | n/a | 24.00 |
| ARID1B | 6 | CAG | 157099403 | 157099455 | 20.67 | 19.67 | n/a | 19.33 |
| ARID1B | 6 | CCG | 157099866 | 157099886 | 7.00 | 7.00 | n/a | 6.67 |
| ARIH1 | 15 | CGG | 72767208 | 72767237 | 12.33 | 12.33 | 11.00 | 9.67 |
| ATN1 | 12 | CAG | 7045880 | 7045938 | 29.33 | 29.33 | 26.67 | 24.33 |
| ATXN1 | 6 | TGC | 16327865 | 16327955 | 37.33 | 37.33 | n/a | 30.00 |
| BCL11B | 14 | CTC | 99641544 | 99641583 | 14.33 | 14.33 | 13.33 | 13.33 |
| BNIP3L | 8 | ACA | 26240684 | 26240700 | 5.67 | 5.67 | 5.67 | 6.33 |
| BRAF | 7 | CGGCGC | 140624404 | 140624425 | 4.33 | 4.33 | n/a | 3.50 |
| BSCL2 | 11 | AGGAGC | 62457909 | 62457946 | 6.67 | 6.67 | 6.33 | 6.17 |
| C2 | 6 | GCC | 31865420 | 31865452 | 11.00 | 11.00 | 11.00 | 10.67 |
| CACNA1D | 3 | GGAAGA | 53766846 | 53766872 | 4.50 | 4.50 | 4.50 | 4.33 |
| CBL | 11 | CAC | 119077233 | 119077254 | 7.33 | 8.33 | 7.33 | 8.00 |
| CBL | 11 | ATG | 119149356 | 119149374 | 6.33 | 7.33 | 6.33 | 7.00 |
| CCND1 | 11 | GAG | 69465976 | 69466003 | 9.33 | 9.33 | 9.33 | 9.33 |
| CD55 | 1 | GCT | 207495170 | 207495188 | 6.33 | 6.33 | 6.33 | 6.00 |
| CD59 | 11 | AGGGCC | 33722001 | 33722022 | 4.17 | 4.17 | 4.33 | 3.50 |
| CDK4 | 12 | GCC | 58149548 | 58149574 | 9.00 | 9.00 | n/a | 9.00 |
| CDT1 | 16 | CGCC | 88870389 | 88870408 | 5.00 | 5.00 | 5.00 | 4.75 |
| CEBPA | 19 | CGC | 33793008 | 33793026 | 7.67 | 7.67 | n/a | 6.00 |
| CEP164 | 11 | CCT | 117280716 | 117280736 | 8.00 | 7.00 | 7.00 | 7.00 |
| CLIP1 | 12 | CT | 122817650 | 122817673 | 12.00 | 12.00 | 12.00 | 12.00 |
| CNGB1 | 16 | CCT | 57983264 | 57983309 | 15.33 | 15.33 | 15.33 | 16.00 |
| CNNM2 | 10 | GCT | 104678378 | 104678411 | 11.33 | 11.33 | 11.33 | 11.00 |
| CNNM2 | 10 | TCC | 104849436 | 104849452 | 5.67 | 5.67 | 5.67 | 5.67 |
| CNPY3 | 6 | TGC | 42897358 | 42897384 | 9.00 | 9.00 | n/a | 9.00 |
| COL1A1 | 17 | CT | 48266555 | 48266569 | 7.50 | 7.50 | 7.50 | 16.00 |
| CTHRC1 | 8 | CTG | 104383933 | 104383957 | 8.33 | 8.33 | 8.33 | 8.33 |
| CTSA | 20 | CTG | 44520238 | 44520263 | 8.67 | 9.67 | 8.67 | 9.33 |
| CYP11B2 | 8 | GCA | 143993951 | 143993965 | 5.00 | 5.00 | n/a | 6.00 |
| DLEC1 | 3 | GCCCCA | 38163910 | 38163937 | 4.67 | 4.67 | 4.67 | 4.50 |
| DMPK | 19 | CGC | 46272031 | 46272052 | 7.33 | 7.33 | 7.33 | 7.00 |
| DPP9 | 19 | AG | 4682879 | 4682893 | 7.50 | 7.50 | 7.50 | 8.00 |
| DUOX2 | 15 | TCCACC | 45393995 | 45394013 | 3.17 | 3.17 | 3.17 | 3.17 |
| EOMES | 3 | GCG | 27763406 | 27763439 | 13.33 | 13.33 | 13.33 | 13.00 |
| ERCC1 | 19 | GAA | 45911859 | 45911873 | 5.00 | 5.00 | 5.00 | 5.00 |
| ERCC1 | 19 | AAG | 45912490 | 45912509 | 6.67 | 6.67 | 6.67 | 16.33 |
| ERCC3 | 2 | TCT | 128046944 | 128046960 | 5.67 | 5.67 | 5.67 | 5.33 |
| ESCO2 | 8 | TCA | 27668477 | 27668492 | 5.33 | 5.33 | 5.33 | 5.00 |
| FANCM | 14 | CT | 45642288 | 45642299 | 6.00 | 6.00 | 6.00 | 6.00 |
| FASLG | 1 | CCA | 172628474 | 172628508 | 11.67 | 11.67 | 11.67 | 11.33 |
| FBXW4 | 10 | CCT | 103454358 | 103454380 | 8.67 | 8.67 | 9.67 | 8.33 |
| FBXW7 | 4 | CTC | 153332605 | 153332628 | 8.00 | 8.00 | 8.00 | 7.67 |
| FERMT3 | 11 | AGA | 63978566 | 63978587 | 7.33 | 7.33 | 7.33 | 7.00 |
| FIP1L1 | 4 | AG | 54319248 | 54319261 | 7.00 | 7.00 | 7.00 | 10.50 |
| FIP1L1 | 4 | CGG | 54966948 | 54966996 | 16.33 | 16.33 | 16.33 | 16.00 |
| FMN2 | 1 | GCA | 240256024 | 240256085 | 20.67 | 20.67 | 20.67 | 20.33 |
| FOXC2 | 16 | CCA | 86602099 | 86602130 | 10.67 | 10.67 | 10.67 | 10.33 |
| FOXE1 | 9 | GCC | 100616214 | 100616232 | 6.33 | 6.33 | 6.33 | 6.00 |
| FOXF1 | 16 | CGG | 86544211 | 86544249 | 14.00 | 13.00 | 14.00 | 12.67 |
| FOXF1 | 16 | GGC | 86544851 | 86544878 | 9.33 | 9.33 | 9.33 | 9.00 |
| FUCA2 | 6 | GCA | 143832708 | 143832734 | 11.00 | 11.00 | 11.00 | 10.67 |
| FUS | 16 | GGC | 31196403 | 31196422 | 10.00 | 8.00 | 6.67 | 12.33 |
| GJC2 | 1 | GAG | 228345913 | 228345933 | 7.00 | 7.00 | 8.67 | 6.67 |
| GJC2 | 1 | CCCCCG | 228346369 | 228346410 | 7.00 | 7.00 | n/a | 6.83 |
| GNAS | 20 | CGAGAC | 57415488 | 57415555 | 11.33 | 11.33 | 11.33 | 11.17 |
| GP6 | 19 | CAGA | 55526092 | 55526121 | 8.50 | 8.50 | 8.50 | 8.25 |
| GPR179 | 17 | CAG | 36490704 | 36490721 | 6.00 | 6.00 | 6.00 | 14.33 |
| GPX1 | 3 | CCCCG | 49395430 | 49395452 | 4.60 | 4.60 | 4.60 | 4.60 |
| GRIN1 | 9 | GGC | 140063408 | 140063434 | 10.00 | 10.00 | n/a | 8.67 |
| HBG2 | 11 | GAA | 5655957 | 5655975 | 7.00 | 7.00 | 6.33 | 7.00 |
| HFE2 | 1 | GAG | 145415369 | 145415387 | 6.33 | 6.33 | 6.33 | 7.00 |
| HLX | 1 | CAG | 221053581 | 221053611 | 15.33 | 10.33 | 10.33 | 11.00 |
| HNF1A | 12 | TCAT | 121434631 | 121434646 | 7.00 | 7.00 | 6.00 | 6.75 |
| HOXA11 | 7 | GCC | 27224167 | 27224183 | 5.67 | 5.67 | 6.00 | 5.33 |
| IGF2R | 6 | CGCC | 160390319 | 160390336 | 4.50 | 4.50 | 4.75 | 4.25 |
| IL27 | 16 | TCC | 28511172 | 28511215 | 16.67 | 14.67 | 14.67 | 14.33 |
| IL4R | 16 | GAG | 27373787 | 27373802 | 5.33 | 5.33 | 5.33 | 5.33 |
| IRF2BP2 | 1 | GGC | 234744786 | 234744809 | 8.00 | 8.00 | 10.00 | 8.00 |
| IRF2BP2 | 1 | GCT | 234744946 | 234744962 | 6.67 | 6.67 | n/a | 6.33 |
| ITGAX | 16 | GAGG | 31368453 | 31368468 | 4.00 | 4.00 | 4.00 | 4.00 |
| KCTD1 | 18 | TCC | 24128310 | 24128338 | 9.67 | 9.67 | 9.67 | 9.33 |
| KCTD1 | 18 | CGG | 24128425 | 24128440 | 5.33 | 5.33 | 5.33 | 5.00 |
| KDM2A | 11 | GAG | 67018067 | 67018098 | 10.67 | 10.67 | 10.67 | 10.33 |
| KDM2B | 12 | CCT | 121947732 | 121947758 | 9.00 | 9.00 | 9.00 | 9.00 |
| KIF1A | 2 | TCC | 241696826 | 241696873 | 17.00 | 17.00 | 17.00 | 16.67 |
| KIF1A | 2 | CTC | 241702149 | 241702163 | 5.33 | 5.33 | 5.00 | 5.00 |
| KIF7 | 15 | CTC | 90189153 | 90189176 | 9.00 | 9.00 | 8.00 | 7.67 |
| KISS1R | 19 | CGC | 920573 | 920600 | 9.33 | 9.33 | 9.33 | 9.00 |
| KITLG | 12 | CT | 88898988 | 88899003 | 8.00 | 8.00 | 8.00 | 7.50 |
| KMT2A | 11 | TCGTCT | 118307495 | 118307517 | 3.83 | 3.83 | 3.83 | 3.67 |
| KRT1 | 12 | GCC | 53069121 | 53069135 | 5.00 | 5.00 | 5.00 | 9.67 |
| KRT1 | 12 | GCC | 53069268 | 53069282 | 5.00 | 5.00 | 5.00 | 5.00 |
| KRT1 | 12 | CCA | 53073735 | 53073762 | 10.33 | 10.33 | 9.33 | 17.00 |
| KRT1 | 12 | CCA | 53073810 | 53073863 | 19.67 | 19.67 | 19.67 | 25.67 |
| KRT10 | 17 | CTC | 38978747 | 38978789 | 15.33 | 14.33 | 14.33 | 15.00 |
| LDB3 | 10 | TGTT | 88492254 | 88492275 | 5.50 | 5.50 | 5.50 | 6.25 |
| LFNG | 7 | GGC | 2559669 | 2559684 | 5.33 | 5.33 | 5.33 | 5.00 |
| LHCGR | 2 | GCA | 48982759 | 48982790 | 12.67 | 12.67 | 12.67 | 12.33 |
| LILRA6 | 19 | GGA | 54723031 | 54723054 | 14.00 | 14.00 | 9.00 | 9.67 |
| LMF1 | 16 | GCG | 1032188 | 1032205 | 6.00 | 6.00 | 6.00 | 5.67 |
| LMNA | 1 | GGC | 156051536 | 156051570 | 11.67 | 11.67 | n/a | 11.33 |
| LURAP1L | 9 | GGC | 12775862 | 12775879 | 9.00 | 9.00 | 9.00 | 8.67 |
| MAN1B1 | 9 | CCG | 139981569 | 139981585 | 5.67 | 5.67 | 5.67 | 7.33 |
| MESP2 | 15 | AGGGGC | 90320121 | 90320204 | 14.50 | 14.50 | 12.50 | 13.83 |
| MGME1 | 20 | GCC | 17949125 | 17949142 | 7.00 | 7.00 | 6.00 | 6.33 |
| MMP20 | 11 | TTTTCC | 102449844 | 102449865 | 3.67 | 3.67 | 3.67 | 4.17 |
| MSH5 | 6 | GAG | 31708352 | 31708381 | 10.00 | 10.00 | 10.00 | 14.67 |
| MTHFR | 1 | TGC | 11866344 | 11866373 | 10.00 | 10.00 | 10.00 | 9.67 |
| MUC5B | 11 | CCA | 1262966 | 1262982 | 5.67 | 5.67 | 5.67 | 5.67 |
| MX1 | 21 | GAA | 42824689 | 42824708 | 6.67 | 6.67 | 6.67 | 6.33 |
| NAGLU | 17 | GGC | 40688492 | 40688509 | 7.00 | 7.00 | 6.00 | 5.67 |
| NBEAL2 | 3 | GCA | 47043575 | 47043602 | 10.67 | 10.67 | 9.33 | 12.00 |
| NKTR | 3 | GAG | 42680269 | 42680283 | 5.00 | 5.00 | 5.00 | 5.00 |
| NKX2-1 | 14 | CCG | 36986880 | 36986901 | 10.33 | 7.33 | 7.33 | 7.33 |
| NOTCH1 | 9 | GTG | 139390945 | 139390960 | 5.33 | 5.33 | 5.33 | 7.00 |
| NPM1 | 5 | TGA | 170819938 | 170819979 | 14.00 | 14.00 | 14.00 | 14.00 |
| NPM1 | 5 | TGA | 170827157 | 170827183 | 9.67 | 9.67 | 9.00 | 8.67 |
| PAH | 12 | CAGCCC | 103352065 | 103352086 | 4.67 | 4.67 | 3.67 | 3.50 |
| PBX1 | 1 | CGG | 164761842 | 164761869 | 9.33 | 9.33 | n/a | 10.00 |
| PCSK9 | 1 | CTG | 55505553 | 55505575 | 9.67 | 8.67 | 8.67 | 8.33 |
| PDX1 | 13 | GCC | 28494391 | 28494417 | 9.00 | 9.00 | 9.00 | 9.00 |
| PIEZO1 | 16 | CCTGCT | 88789667 | 88789684 | 4.50 | 4.50 | 4.00 | 2.83 |
| PLEC | 8 | CCAGCT | 144998252 | 144998271 | 4.17 | 4.17 | 3.33 | 3.17 |
| PML | 15 | CCAGCC | 74287252 | 74287275 | 4.00 | 4.00 | n/a | 4.83 |
| PNPLA2 | 11 | TCCC | 822001 | 822015 | 3.75 | 3.75 | 3.75 | 3.50 |
| POLG | 15 | GCT | 89876820 | 89876860 | 18.67 | 15.67 | 14.67 | 15.33 |
| POMC | 2 | GCT | 25384460 | 25384481 | 13.33 | 10.33 | 10.33 | 10.00 |
| PRKCSH | 19 | GAG | 11558341 | 11558405 | 23.33 | 23.33 | 23.67 | 22.33 |
| PTF1A | 10 | CGG | 23481741 | 23481756 | 5.33 | 5.33 | 5.33 | 5.00 |
| PTF1A | 10 | GCG | 23481889 | 23481921 | 11.00 | 11.00 | 11.00 | 10.67 |
| RAI1 | 17 | CAG | 17697094 | 17697134 | 17.67 | 17.67 | 13.67 | 14.33 |
| RAI1 | 17 | GCA | 17699993 | 17700010 | 6.00 | 6.00 | 6.00 | 6.33 |
| RB1 | 13 | GCC | 48878076 | 48878101 | 8.67 | 8.67 | 8.67 | 8.67 |
| RECQL4 | 8 | GGTGCA | 145738411 | 145738430 | 4.33 | 4.33 | 3.33 | 3.17 |
| RLTPR | 16 | CCGCG | 67682307 | 67682326 | 4.00 | 4.00 | 4.00 | 3.80 |
| RSPH4A | 6 | GAA | 116950769 | 116950800 | 10.67 | 10.67 | 10.67 | 11.00 |
| RYR1 | 19 | GAG | 38979895 | 38979962 | 24.33 | 24.33 | 22.67 | 23.67 |
| SCN9A | 2 | TTTC | 167143002 | 167143028 | 6.75 | 6.75 | 6.75 | 6.50 |
| SERPING1 | 11 | CTG | 57365774 | 57365788 | 5.00 | 5.00 | 5.00 | 5.00 |
| SETX | 9 | TCA | 135203911 | 135203928 | 6.00 | 6.00 | 6.00 | 6.67 |
| SFTPB | 2 | GCA | 85895264 | 85895280 | 7.67 | 7.67 | 5.67 | 6.00 |
| SFTPC | 8 | GTG | 22020159 | 22020175 | 6.67 | 5.67 | 5.67 | 6.33 |
| SHH | 7 | GCC | 155595750 | 155595781 | 10.67 | 10.67 | 11.67 | 10.33 |
| SHISA6 | 17 | CTG | 11144758 | 11144774 | 5.67 | 5.67 | n/a | 5.33 |
| SIX3 | 2 | GGC | 45169361 | 45169395 | 12.67 | 12.67 | n/a | 11.33 |
| SLC19A2 | 1 | GCC | 169454958 | 169454980 | 8.67 | 7.67 | 7.67 | 7.33 |
| SLC24A1 | 15 | GAG | 65943068 | 65943157 | 30.00 | 30.00 | 31.67 | 29.67 |
| SMPD1 | 11 | CTGGCG | 6411931 | 6411971 | 8.83 | 8.83 | 6.83 | 6.67 |
| SOX10 | 22 | CCG | 38379676 | 38379690 | 5.00 | 5.00 | n/a | 4.67 |
| SOX11 | 2 | GCG | 5833277 | 5833296 | 6.67 | 6.67 | 6.67 | 6.33 |
| SOX11 | 2 | GAC | 5833526 | 5833554 | 9.67 | 9.67 | 9.67 | 11.33 |
| SOX11 | 2 | CAG | 5833885 | 5833915 | 14.33 | 14.33 | 14.33 | 10.00 |
| SOX17 | 8 | CACCAG | 55372259 | 55372287 | 5.83 | 5.83 | 4.83 | 5.67 |
| SOX18 | 20 | GCGG | 62680707 | 62680744 | 9.50 | 9.50 | n/a | 9.25 |
| SPAG1 | 8 | GCG | 101225491 | 101225515 | 11.67 | 11.67 | 8.33 | 8.67 |
| SPHK2 | 19 | GGCTGG | 49132424 | 49132449 | 4.33 | 4.33 | 4.33 | 7.33 |
| SPINK5 | 5 | AGA | 147480924 | 147480948 | 8.33 | 8.33 | 8.33 | 9.33 |
| SPTB | 14 | TCTCC | 65219002 | 65219029 | 6.60 | 6.60 | 5.60 | 6.40 |
| STIM1 | 11 | CAC | 4107924 | 4107960 | 12.33 | 12.33 | 12.33 | 13.33 |
| TBX1 | 22 | CGC | 19748552 | 19748600 | 16.33 | 16.33 | n/a | n/a |
| TBX1 | 22 | CAC | 19754259 | 19754274 | 7.00 | 7.00 | n/a | 5.33 |
| TBX1 | 22 | GCC | 19754286 | 19754330 | 15.00 | 15.00 | n/a | 14.67 |
| TET3 | 2 | GGA | 74328808 | 74328826 | 6.33 | 6.33 | 6.33 | 7.00 |
| TGFB1 | 19 | CAG | 41858902 | 41858935 | 11.33 | 11.33 | 11.33 | 12.00 |
| TGM1 | 14 | TCTGGC | 24731464 | 24731486 | 4.83 | 3.83 | 3.83 | 3.67 |
| TMEM43 | 3 | TCCT | 14177451 | 14177465 | 3.75 | 3.75 | 3.75 | 3.50 |
| TTN | 2 | TCT | 179544686 | 179544700 | 6.00 | 6.00 | 6.00 | 5.67 |
| TULP1 | 6 | CCT | 35477045 | 35477073 | 9.67 | 9.67 | 9.67 | 9.67 |
| UNC93B1 | 11 | CTC | 67771218 | 67771232 | 5.00 | 5.00 | 5.00 | 4.67 |
| WDR81 | 17 | GAG | 1631749 | 1631776 | 11.00 | 11.00 | 9.33 | 9.00 |
| WT1 | 11 | GGC | 32456485 | 32456524 | 13.33 | 13.33 | 13.33 | 13.00 |
| XKR6 | 8 | GCC | 11058714 | 11058740 | 9.00 | 9.00 | 9.00 | 9.00 |
| ZIC2 | 13 | GCG | 100634394 | 100634410 | 5.67 | 5.67 | 5.67 | 5.67 |
| ZIC2 | 13 | GCG | 100637703 | 100637748 | 15.33 | 15.33 | 17.00 | 15.00 |
| ZIC2 | 13 | GGC | 100637805 | 100637861 | 19.00 | 19.00 | n/a | 18.67 |
| ZMIZ1 | 10 | CTC | 81070787 | 81070803 | 6.67 | 5.67 | 5.67 | 5.33 |
